# Cleaner wrasse failed in early testing stages of both visual and spatial Working Memory paradigms

**DOI:** 10.1101/2025.01.16.633362

**Authors:** Leonore Bonin, Héctor M. Manrique, Redouan Bshary

## Abstract

Working memory (WM), an attention-based short-term storage system responsible for the manipulation and integration of past knowledge with present information for goal-directed behavior, is a key executive function and a principal predictor of general intelligence. As WM has not been a major research topic in animal behavior, we first summarize key ideas from the social sciences for interested colleagues. Given that past methodological inconsistencies have led to mixed results and conclusions across various species, we designed experiments that incorporate the critical components of WM, facilitating cross-species comparisons and accounting for potential ecological influences. We present such experiments on WM in an ectothermic vertebrate, the cleaner wrasse (*Labroides dimidiatus*), which faces environmental challenges potentially requiring complex cognitive adaptations. Overcoming several experimental challenges, we consistently obtained negative results across multiple experimental paradigms. Our negative results using experiments specifically designed to test WM call into question previous studies that used other experiments and reported positive WM findings in other fish species. If the negative results in specific WM tests were to be confirmed in other ectotherm vertebrates, the absence of WM may turn out to be a key factor underlying the significant encephalization gap between ectotherm and endotherm vertebrate species.

## Introduction

Among vertebrates, brain size relative to body size exhibits considerable variation (Jerison, 1969). On average, endotherms possess brains that are approximately ten times larger than those of ectotherms when corrected for body size (Jerison, 1969). Given the high energetic demands of brain tissue (Fonseca-Azevedo & Herculano-Houzel, 2012; Sukhum et al., 2016), an important question arises: what advantages do larger brains confer to endotherms? One hypothesis is that endotherms possess a more extensive cognitive toolkit compared to ectotherms. While both groups exhibit basic learning strategies that allow them to form new stimulus-response associations (e.g., Pavlovian conditioning, Pavlov, 2010), and/or learn new adaptive responses following the presentation of a reward (Thorndike, 1927), these strategies do not require flexible manipulation of mental objects. Whether ectotherms are capable of “complex cognition”, e.g., cognitive abilities involving the executive functions of the brain (EFs, also called executive or cognitive control) (Knauff & Wolf, 2010), remains poorly understood and warrants further exploration.

A complementary hypothesis regarding the differences in brain size between endotherms and ectotherms relates to the sophistication of these EFs. These higher-level cognitive skills are central to complex cognition, as they involve building intricate interactions among mental processes of varying complexity (Diamond, 2013). The three primary EFs—inhibition, cognitive flexibility, and working memory—“are recruited when it would be ill-advised, insufficient, or impossible to go on autopilot or rely on instinct or intuition, such as when presented with novel, unanticipated challenges” (Diamond, 2020). Although research on EFs in ectotherms is scarce, some studies suggest that fish, for example, perform well in tasks assessing cognitive flexibility as shown in reversal learning (Aellen et al., 2022; Parker et al., 2012) – where performance correlates positively with brain size in primates (Deaner et al., 2007) – and demonstrated proficiency in inhibitory control tasks, such as delayed gratification paradigms (Aellen et al., 2021) and detour tasks (Lucon-Xiccato & Bisazza, 2017; MacLean et al., 2014).

In this study, we investigated the role that working memory (WM), a critical EF, could play in ectotherms’ cognition. There is a vast literature on WM in the social sciences. Research on WM includes comparative developmental psychology, i.e. comparisons between children and great apes. While definitions of WM may have become more crystallized in recent years, there are rather diverse ideas on how to test for WM (Manrique et al. 2024). As we consider such aspects critical for future research on WM in animals, we first provide a brief summary of definitions and methodological considerations as discussed in detail by Read et al. (2022) and Manrique et al. (2024).

### WM definition and importance

WM is among the most well-studied EFs in humans, with extensive research dating back more than 80 years to Miller et al. (1960), who referred to “the memory we use for the execution of plans as a kind of quick access working memory” (p. 65). While numerous models of WM have been proposed, we focus on the Embedded-Processes model of Cowan (1988, 1995), which has undergone recent refinements (Cowan et al., 2024), and it is not reliant on language. We redirect readers to appropriate literature for descriptions and debates about other models (e.g. Baddeley & Hitch, 1974; Chai et al., 2018; Conway et al., 2003; Cowan, 1988, 1995; Cowan et al., 2024; Fuster & Bressler, 2012; Logie et al., 2021; Morrison, 2005). The Embedded-Processes model consists of a focus of attention (FoA) within a temporarily activated portion of the long-term memory (aLTM). Cowan et al. (2024) defined the FoA as a “coherent representation of several separate items or ideas guiding current thoughts and actions” and the aLTM as “the part of memory from which information is in a heightened state of accessibility”. Hence, in this model, a new stimulus can enter either consciously with the intervention of the central executive, or unconsciously into the aLTM. Part of the information related to that stimulus enters directly into the FoA and reaches awareness. Thanks to the central executive, some knowledge stored in the LTM can be brought into the FoA to “guide focus”. By combining them, new “chunks” are created and can move from the FoA to the aLTM, and finally to the LTM where this new acquired knowledge will remain stored. In line with this model, Manrique et al. (2024) defined WM as a “brain system that provides us with temporary short-term storage and management of perceptual or other information (…), which we need for efficiently (…) carrying out, and updating, such complex cognitive tasks as mental reading, reasoning, forecasting, manipulation, (…)”. This is the definition we followed in this study. The manipulative aspect of WM distinguishes it from short-term memory.

WM is of potential importance because it has been linked to many cognitive abilities. For example, training WM can improve crystallized intelligence (Alloway & Alloway, 2009). WM is also a major predictor of *g* in mice (Kolata et al., 2005)—a species representing an endotherm with rather average brain size-body size relationship (Tartarelli & Bisconti, 2006)—and in humans (discussed in Conway et al., 2003). WM is also correlated with numerous other cognitive and perceptual abilities in humans (Ackerman et al., 2002; Engle, 2010; Unsworth et al., 2015), and the prospective aspect of WM is key to efficient tool crafting (Manrique & Walker, 2017). At present, the only experiment that has tested the general intelligence factor *g* in fish yielded negative results (Aellen et al., 2022), suggesting that their WM might indeed be more primitive and/or not used in a domain-general way. Alternatively, ectotherms might have WM capacities similar to endotherms, but these capacities could be modular and only elicited under ecologically relevant circumstances.

### How to test for WM?

There are not only multiple definitions of WM, but also different opinions on how to test it. Cowan et al. (2024) argue that “Discrepancies between methods or definitions often underlie discrepancies between results rather than unreliability of evidence”, highlighting the importance of producing a clear and common methodology. Testing WM in animals presents distinct challenges, especially in the absence of language. Tasks commonly used in human studies, such as verbal WM tasks (Hu et al., 2019), are not feasible for animals.

While non-verbal tasks have yielded interesting results, caution must be exercised due to the limitations of these methodologies, including the lack of a manipulative or problem-solving aspect. This problem is readily apparent in some spatial tasks. For example, in zebrafish, free exploration of a Y-maze and spontaneous alternation has been taken as evidence of WM (Fontana et al., 2021). Similarly, testing for the Piagetian stage 4 of object permanence abilities has also been taken as evidence of WM in cichlids (Guadagno & Triki, 2024) and in dogs (with time delay, Fiset et al., 2003) even though it is not evident that recovering the object involves the mental manipulation of temporarily stored information. This criticism does not apply to higher stages of object permanence (i.e., Piagetian stages 5 and 6) where the position of the object must be subsequently updated, thus imposing higher demands on WM. These latter stages have already been used to test for WM in corvids and psittacids (Pepperberg & Funk, 1990; Zucca et al., 2007).

Zebrafish performance on a delayed-match-to-sample task has been taken as evidence of WM (Bloch et al., 2019), and the same paradigm has also been applied to honeybees (Zhang et al., 2005) yielding negative results. However, these types of tasks rely heavily on visual perceptual information and could be seen as “recognition tasks” (Bachevalier & Mishkin, 1986; Zola et al., 2000). If so, they could be solved by relying on non-conscious processes, such as familiarity. Whether recognition of a previously presented stimulus is a reliable measure of WM is debatable (Yonelinas, 2002), and it would be crucial to test whether success in such a task depends on recollection or simply familiarity.

Recent studies have also identified methodological limitations when testing WM in nonhuman primates, such in the “limited-hold memory task” applied to measure memory of chimpanzees (Inoue & Matsuzawa, 2007). This task may instead rely on some imagery mechanisms that do not tap into WM (Abdi et al., 2022; Manrique & Walker, 2017; S. Miller & Peacock, 1982; Özbaydar, 2012; Read et al., 2022). Many other methodological and conceptual limitations have been identified in relation to the measurement of WM in great apes, which are discussed extensively in Read et al. (2022) and Manrique et al. (2024).

Another important consideration when testing WM involves selecting the appropriate time interval separating the acquisition from the retrieval of information (Carruthers, 2013). This is not trivial, as WM is connected to other short-term storages, but also, critically, to the activation of portions of information stored in long-term memory. Further involvement of long-term memory in the resolution of a task might also depend on whether participants have time to combine the information (Morey & Cowan, 2018). Sensory memory, the non-manipulative short-time storage, is known to last from milliseconds to a maximum of 4 seconds in humans (Darwin et al., 1972; Sperling, 1960). Previous studies often failed to sufficiently explain the specific time interval chosen to elicit WM; measurements can range from 0 sec (e.g., Fiset et al., 2003) to 15 min (Fontana et al., 2021), which can easily lead to confusion and mixed results. In spatial tasks, such as in maze experiments (Kraeuter et al., 2019), a potential shortcoming is that the time intervals used, and the number of items to be remembered, may link performance more to long-term WM (Carruthers, 2013).

The last issue we want to discuss is the involvement of attention, which is notably highlighted in Cowan’s Embedded-Processes model (1988, 1995). Sensitivity to disturbance is a critical characteristic of WM testing (e.g., Angelopoulou & Drigas, 2021; Cowan, 1988; Cowan et al., 2024) that has been largely overlooked in previous publications (e.g., Doré et al., 1996; Lee et al., 2015; Macpherson & Roberts, 2010).

Despite these methodological challenges, cross-species comparisons of WM can provide valuable insights. WM may vary quantitatively rather than qualitatively among species. For example, humans can maintain 3-5 objects in their FoA, but are also able to almost double this number through exerting executive control over the contents of memory (Manrique et al., 2024; Read et al., 2022). In contrast, other species with lower cognitive capacities may access only one or two objects and may be less capable of combining information to form higher chunks (Cowan, 2022; Cowan et al., 2024).

### The current study

We used the cleaner wrasse *Labroides dimidiatus* (“cleaner” hereafter) for our study as it constitutes a suitable model organism to explore the potential cognitive tool kit of ectotherms. Despite exhibiting an average brain-to-body size ratio for a fish (Chojnacka et al., 2015), cleaners possess a remarkable cognitive tool kit (Bshary & Triki, 2022) that includes basic perspective-taking (McAuliffe et al., 2021), transitive inference (under certain circumstances, Bonin & Bshary, 2023; Hotta et al., 2020), generalized rule learning (Wismer et al., 2016), chaining and configurational learning (Prat et al., 2022; Quiñones et al., 2020), true mirror self-recognition (Kohda et al., 2019, 2022, 2023), and reversal learning (Triki & Bshary, 2021). Particularly relevant to our study is their ability to delay gratification, which is comparable to that of non-human primates (Aellen et al., 2021), despite an apparent lack of the general intelligence factor *g* (Aellen et al., 2022). Furthermore, cleaners remembered the *when* and the *what* after 2.5 - 15 minutes in a foraging task (Salwiczek & Bshary, 2011). Although this may qualify as episodic-like memory (Manrique & Walker, 2017; Pause et al., 2013; Tulving, 1972, 2002), it raises the question about WM as cleaners seem to adjust and update their behavior flexibly in response to previously acquired knowledge. This prompted us to modify the experiment such that solving the *when* and *what* would require WM.

To encompass several aspects of the definitions of WM, we designed experiments that varied in both the level of complexity and the specific context, aiming to provide a comprehensive preliminary assessment of WM in cleaners. We present the results from four experiments. These experiments involve, in part, adjustments from three others that led to cleaners behaving such that we considered any data to be inconclusive. Please see supplementary material for additional methodologies and explanations for experimental designs that we abandoned.

The first experiment we present, referred to as the “windows experiment”, involved placing food items inside Plexiglas rings (windows) attached to a Plexiglas plate, rendering the items invisible when the fish approached the location from the side. We also varied the ecological relevance by using a plain white plate or a fish picture to assess whether this influenced performance. Although this experiment was intended as a training step for a more complex one (see supplementary material), the data provided initial insights into the investigation of WM in cleaners.

The second experiment we present involved a movable arena where cleaners had to make several choices to access and eat all the hidden food items on a compartmentalized plate. In many regards, this was similar to Völter et al.’s (2019) self-ordered search task where chimpanzees had to look for food rewards in different boxes while avoiding repeating a previously searched box, relying either on the spatial location that the boxes occupied in previous trials or in their exterior perceptual appearance. Experiments 1 and 2 have in common that we investigated WM with respect to the spatial distribution of food items.

In the remaining two experiments, we investigated the potential presence of WM in a social context. In experiment 3, we used a methodology similar to Salwiczek & Bshary (2011), first letting subjects eat a single item on the back of one plate, removing the plate, and then reintroducing the same (now empty) plate alongside a second one that still had a food item on its back. In experiment 4, we modified the methodology so that the two plates were always visible to cleaners, addressing the possibility that negative results in experiment 3 may have been due to subjects having misunderstood the task by interpreting the reintroduced plate as a new object, despite its identical color.

To ensure that our experiments 2-4 did not fall into the domain of sensory memory (Darwin et al., 1972; Sperling, 1960), we set a minimum delay of 5 seconds separating potential encoding and retrieval. We set a maximum delay of 10 seconds to limit the possibility that differences in information rehearsing abilities could overshadow differences in WM (Morey & Cowan, 2018) and avoid measuring abilities strictly related to long-term memory (LTM), as previous research suggests that in humans, information can transfer into LTM within 15 to 18 seconds (Atkinson & Shiffrin, 1968; WM and LTM discussed in Rhodes & Cowan, 2018 for example). Delay times could be extended in future studies to investigate the influence of delay duration in greater detail.

Given the apparent lack of a general intelligence factor *g* in cleaners, we anticipated predominantly negative results, particularly in the more complex experiments. One of our main objectives was to assess whether the correlation between WM and *g* observed in other species holds for cleaners, with the absence of *g* implying limited WM capacity. However, cleaners frequently encounter multiple-choice scenarios in their natural environment that require the integration of previous knowledge into current decision-making. Thus, we also expected potential positive results in experiments of lower complexity. As demonstrated in rats (Bratch et al., 2016), cleaners may possess different WM sub-processes, allowing for WM measurements across diverse contexts.

## General methods

The study was conducted during two field trips in 2021 and 2022 to Mo’orea, French Polynesia. We caught adult cleaner fish in non-protected reefs near Cook’s Bay. We led cleaners into a barrier net (mesh size: 0.5cm) using hand nets. Then, we placed them into zip-loc bags for transport and renewed the water after 45 minutes to ensure that it contained enough oxygen. Fish were housed at the Gump Research Station in white opaque plastic aquaria, 70cm (length) x 52cm (width) x 45 cm (height), with a water height of around 30-35cm. The water came directly from the ocean and was in an open circuit, closed during experiments only. Cleaners were first habituated living in the aquaria, and not tested until 8-15 days post-capture, during which time they learned to feed on plates and became accustomed to the basic material and experimenters.

A camera placed on the experimenter’s forehead recorded each trial.

All analyses were run with R software v. 4.3.1 (R Core Team, 2022). For each model, we conducted an analysis of variance to compare two versions: one with the fish identity as a random factor for both the intercept and the slope with respect to time (trial number) and the other only with the fish identity as a random factor. We always kept the simpler versions as those were not significantly different, and they had slightly smaller AIC.

The French Polynesian authorities approved the catching and the manipulation of cleaners (11626/VP/DRM). After the experiments, we released the cleaners at their site of capture.

All datasets and R scripts are accessible on Figshare (https://figshare.com/projects/Cleaner_wrasse_failed_in_early_testing_stages_of_both_visual_ and_spatial_working_memory_paradigms/231434).

## Specific methods and results

For clarity, we present the methods and results together for each experiment.

### 1. Experiment 1, windows experiment

#### Methods

Nine adult females, caught between 18.-20.9.21, were tested on four days from 7-10.10.21. These fish had previously taken part in abandoned experimental designs described in the supplements. For the experiment, we used two different plates with black spots marking the locations of four food items: (1) a plain white square (hereafter referred to as “square”, 20cm x 20cm, Fig. 1a) and (2) a laminated picture of a butterflyfish (*Chaetodon ornatissimus*, hereafter referred to as “fish”, 20cm length x 14cm height, Fig. 1b). Locations of items were up, right, down and left, which corresponded to specific areas on the fish plate (e.g., dorsal fin, gills, anal fin and caudal fin) to mimic natural ecologically meaningful spots. We applied two conditions to both plates: (i) ‘basic’ where four food items are always visible, and (ii) ‘window’ where the four food items are placed inside Plexiglas wells of 1.5 x 1.5cm and hence become invisible unless faced, Fig. 1a-b).

**Figure 1:**
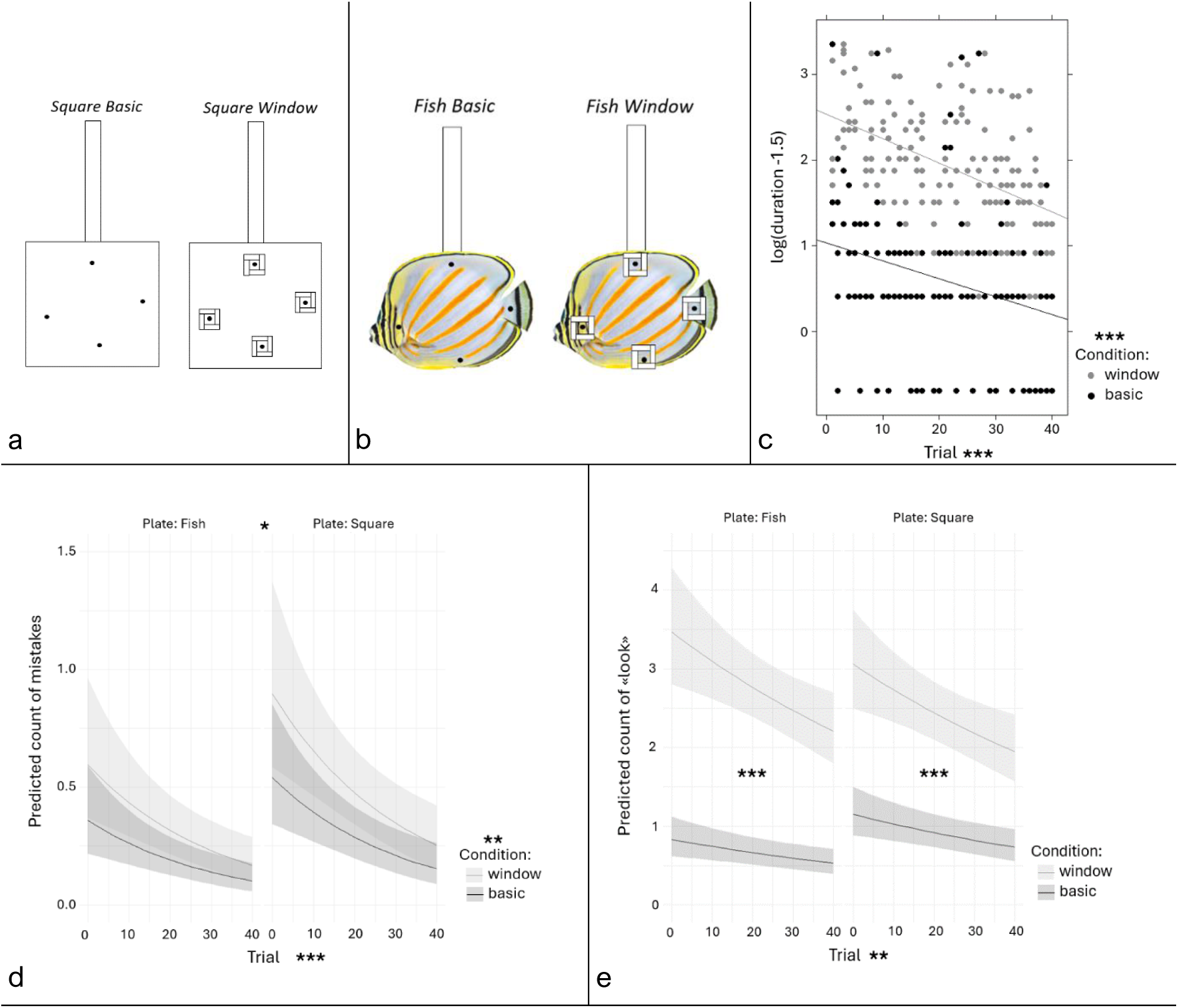
Windows experiments methods (a, b) and results (c, d, e) **a. White square plate and b. butterflyfish plates.** We used both types of plates in basic and window conditions. The black spots held mashed prawn items. Each window was 0.5cm deep. **c. Effect of time and condition on the duration to eat the four items.** The y-axis represents the log-transformed duration, and the x-axis shows the trial number. The color code indicates the condition. The stars indicate the significance of the associated variable. Significance codes: 0 ‘***’ 0.001 ‘**’ 0.01 ‘*’ 0.05 ‘.’ 0.1 ‘ ‘ 1. **d. Effects of plate, condition, and time on the number of mistakes.** The y-axis is the predicted mistakes count, and the x-axis displays the trial number. Each panel represents a plate, and the color indicates the condition. The stars indicate the significance of the associated variable. Significance codes: 0 ‘***’ 0.001 ‘**’ 0.01 ‘*’ 0.05 ‘.’ 0.1 ‘ ‘ 1. **e. Effect of plate, condition, and time on the number of times cleaner looked around.** The y-axis is the predicted count of times cleaners looked around (*i.e.*, opportunities to see the items again), and the x-axis is the trial number. Each panel represents a plate, and the color code indicates the condition. The significance of the difference between both condition lines is given on each panel, and the significance of the covariate “Trial” is shown next to the x-axis label. Significance codes: 0 ‘***’ 0.001 ‘**’ 0.01 ‘*’ 0.05 ‘.’ 0.1 ‘ ‘ 1.

We conducted ten trials per fish per day for four consecutive days. Of the nine cleaners, four initially performed 20 trials using the fish plate, while five began with the square plate. The condition (basic or window) for each trial was counterbalanced, ensuring that no more than two consecutive trials featured the same condition. Each condition was tested five times per day.

Each trial followed the same procedure: the cleaner was placed behind a transparent barrier, with the plate inserted on the opposite side of the aquarium. Once the cleaner was released, we recorded the sequence in which it ate the food items, and counted the number of mistakes (i.e. the number of times it picked at an already-eaten location). Based on observations during data collection, we also extracted additional information from video footage, including: the total time taken to eat from all locations (if achieved, hereafter called “duration”), instances of “inspection” (i.e., the cleaner swimming around the plate, sometimes visually examining the locations without picking on the black spot), and instances of “leaving” (when the fish moved at least 5 cm away from the plate, marked by a line at the aquarium’s base). Both inspection and leaving behaviors, which were grouped under the categorical variable “looks”, allowed the fish to re-examine (i.e. look again) the plate, potentially preventing mistakes. These three variables (number of mistakes, duration, and looks) were then used as three distinct response variables to estimate foraging efficiency. Plates were left in the aquarium for a maximum of one minute, but data were only extracted for the first 30 seconds, as eating the four items should occur fairly quickly.

##### Statistical analyses

We categorized our measurements into three response variables: the duration of time (maximum of 30 seconds) taken to eat the food items from all four locations, the number of mistakes (repeated location), and the number of “looks”. We aimed to assess the effects of ecological relevance (represented by the plate: square or fish), visual information (or complexity, represented by the condition: basic or window), and learning (examined through performance trends across trials). We began with the most complex model, including all possible interaction terms, and performed model selection to identify the best fit.

Discarded trials: Trials in which subjects failed to eat items from all locations within 30 seconds were excluded from the analysis. Additionally, one trial on Day 2 (8th trial) was omitted due to a camera malfunction that prevented data extraction, and one trial for Fish E (Day 1, 10th trial) was excluded because the plate was removed prematurely, preventing access to all locations. Duration data: Despite applying various transformations to our duration data, the distribution remained problematic, with none yielding homogenous variances. The best-fitting model was a linear mixed-effects model applied to log-transformed duration data, with a transformation parameter of −1.5. The explanatory variables included the plate, condition, and trial number, along with all interaction terms. The identity of cleaners subjects was treated as a random factor, and heteroscedasticity was modelled between the “basic square” condition and other combinations using the VarFunc class from the *nlme* package (Pinheiro et al., 2007) in R. We simplified the model by retaining only the plate, condition, and trial number as fixed effects.

Mistakes and looks data: As both response variables represented count data, we used generalized linear mixed-effects models with a Poisson distribution. The same methodological approach as used for the duration analysis was applied, leading to similarly simplified models. Models diagnoses and post-hoc analyses: Model diagnostics were performed using the *DHARMa* package (Hartig, 2022) in R, where we examined outliers and checked for model assumptions. We retained all outliers in our analyses as there was no valid justification for their removal. Overdispersion was checked when required. For post-hoc analyses, we employed least-square means methods from the *emmeans* package (Lenth, 2023) in R.

#### Results

Among the 350 trials left for analyses, cleaners did not eat at all locations after 30 seconds in 19 of them. Among these, two were in the basic condition vs. 17 in window condition, and nine were with the fish plate vs. 10 with the square plate. Also, these trials were spread among eight of the nine cleaners (leading to the removal of 1-4 trials from analyses, respectively).

Duration: With the full model (i.e., with all possible interactions), the condition:trial number was only marginally significant (Type II Wald Chisquare tests, condition:trial, Chisq = 2.92, df = 1, p-val = 0.09). We first simplified the model by keeping only that interaction along with the other covariates to see whether the tendency would be confirmed. As it came out non-significant (Type II Wald Chisquare tests, condition:trial: chisq = 2.22, df = 1, p-val = 0.14), we finally removed it as well and kept the simpler model with the three separated covariates.

Cleaners took 2.55 more time to eat the four items in the windows compared to the basic condition (least-square means, basic: predicted mean = 3.31sec, SE = 0.15 and windows: predicted mean = 8.45sec, SE = 0.54, significance from Type II Wald Chisquare tests, condition: chisq = 354.95, df = 1, p < 2e-16, Fig. 1c). There was also improvement of performance over time; cleaners were almost twice as fast to eat the four items on trial 40 compared to the first trial (least-square means, trial 1: predicted mean = 7.43sec, SE = 0.55 and trial 40: predicted mean = 3.69 sec, SE = 0.20, significance from Type II Wald Chisquare tests, trial: chisq = 73.12, df = 1, p-val < 2e-16, Fig. 1c). In contrast, the plate type (fish or square) only showed a tendency (Type II Wald Chisquare tests, chisq = 2.77, df = 1, p-val = 0.096).

Mistakes: None of the interaction terms was significant in the full model (Type II Wald Chisquare tests, chisq max = 2.65, p-val min = 0.10, all df = 1). After simplification, we found that cleaners made around 1.7 times fewer mistakes in basic compared to windows condition (least-square means, basic: predicted mean = 0.23, SE = 0.04 and windows: predicted mean = 0.38, SE = 0.06, significance from Type II Wald Chisquare tests, condition: chisq = 6.84, df = 1, p-val = 0.009, Fig. 1d). Note that ‘mistakes’ include cases in which the fish had not eaten the entire prawn item, and hence ate a small leftover. This explains why we scored mistakes even in the condition in which everything was visible. There was also improvement over time; the number of mistakes was 3.4 times smaller at trial 40 compared to trial 1 (least-square means, trial 1: predicted mean = 0.55, SE = 0.1 and trial 40: predicted mean = 0.16, SE = 0.04, significance from Type II Wald Chisquare tests, trial: chisq = 13.69, df = 1, p-val < 0.001, Fig. 1d).

Finally, cleaners made 1.5 times more mistakes (i.e., repeating a location) in the square compared to the fish plate (least-square means, square: predicted mean = 0.36, SE = 0.05 and fish: predicted mean = 0.24, SE = 0.04, significance from Type II Wald Chisquare tests, plate: chisq = 4.22, df = 1, p-val = 0.04, Fig. 1d).

Looking: In the full model, only the interaction between plate and condition came out significant (Type II Wald Chisquare tests, chisq = 5.15, df = 1, p-val = 0.023). In the simplified model, the interaction between plate and condition remained significant (Type II Wald Chisquare tests, chisq = 5.10, df = 1, p-val = 0.02). The interaction exists because when tested with the fish plate, cleaners looked 4.2 times less in basic than in window condition, while the factor was only 2.6 in the square plate condition (fish: basic: predicted mean = 0.66, SE = 0.09 and windows: predicted mean = 2.73, SE = 0.20, z-ratio = −9.70, p-val < 0.0001; square plate: basic: predicted mean = 0.91, SE = 0.10 and window: predicted mean = 2.41, SE = 0.18, z-ratio = - 7.21, p-val < 0.0001, Fig. 1e).

Overall, performance improved over trials (Type II Wald Chisquare tests, trial: chisq = 9.04, df = 1, p-val = 0.003, Fig. 1e). In the first trial, cleaner fish had to look 1.55 times more often compared to trial 40 (trial 1: predicted mean = 1.77, SE = 0.16 and trial 40: predicted mean = 1.77, SE = 0.11).

### 2. Experiment 2, Working Memory box: the movable arena

#### Methods

We started the experiment with eight males that had been caught between 24. – 28.1.22. Three males did not properly habituate to the experimental design, leaving us with five males. Trials took place between 3. – 17.3.22. subjects had been exposed to mirrors before, which should not have affected their performance in any obvious way.

The experimental design represents a more sophisticated version of the first experiment, testing again the subjects’ ability to remember which food locations they have already visited. At the start of each trial, the tested cleaner entered the arena when the experimenter opened the first door (door 1, Fig. 2a) and started the trial when passing the second door (door 2, Fig. 2a) which led to the center of the arena (35cm length, 20cm width, Fig. 2a). Swimming through, cleaners could access a square plate (20cm x 20cm) that had four round holes with pipes of 2 cm diameter and about 2.5cm length in it (Fig. 2c-d) leading to individual, separated compartments (square of 10cm × 10cm and 7cm deep, Fig. 2a). A piece of latex glove used as curtain prevented the fish from seeing what was behind the hole and, in most cases, prevented cleaners from coming out the compartment through the pipe. In each compartment, there was one mashed prawn item placed on the central side wall (Fig. 2d). Because of the rubber curtain, cleaners had to exit compartments by the individual attending corridors (total length 50cm, height and width 5cm, Fig. 2a) that led to the initial starting point. Only from there could cleaners swim again through the still-open door 2 and choose again between the four holes. We let cleaners repeat such cycles until they had eaten all four food items (details of the procedure provided in Fig. 2a–d). The distances were such that the time interval from consumption of a food item until the next decision was typically within the 5-10s interval that we had aimed for as long as the fish swam at normal speed. We did not quantify time within test trials, but trials in which subjects stopped swimming were not considered. Note that the external walls of the arena were dashed and the corridors were built using blurred see-through Plexiglas so that cleaners could still perceive the presence of walls but light could easily go through the arena. The advantage of the current design over a radial maze is that subjects always come back to the same starting point between choices, making it impossible for a simple swimming routine to explain a potential success.

**Figure 2:**
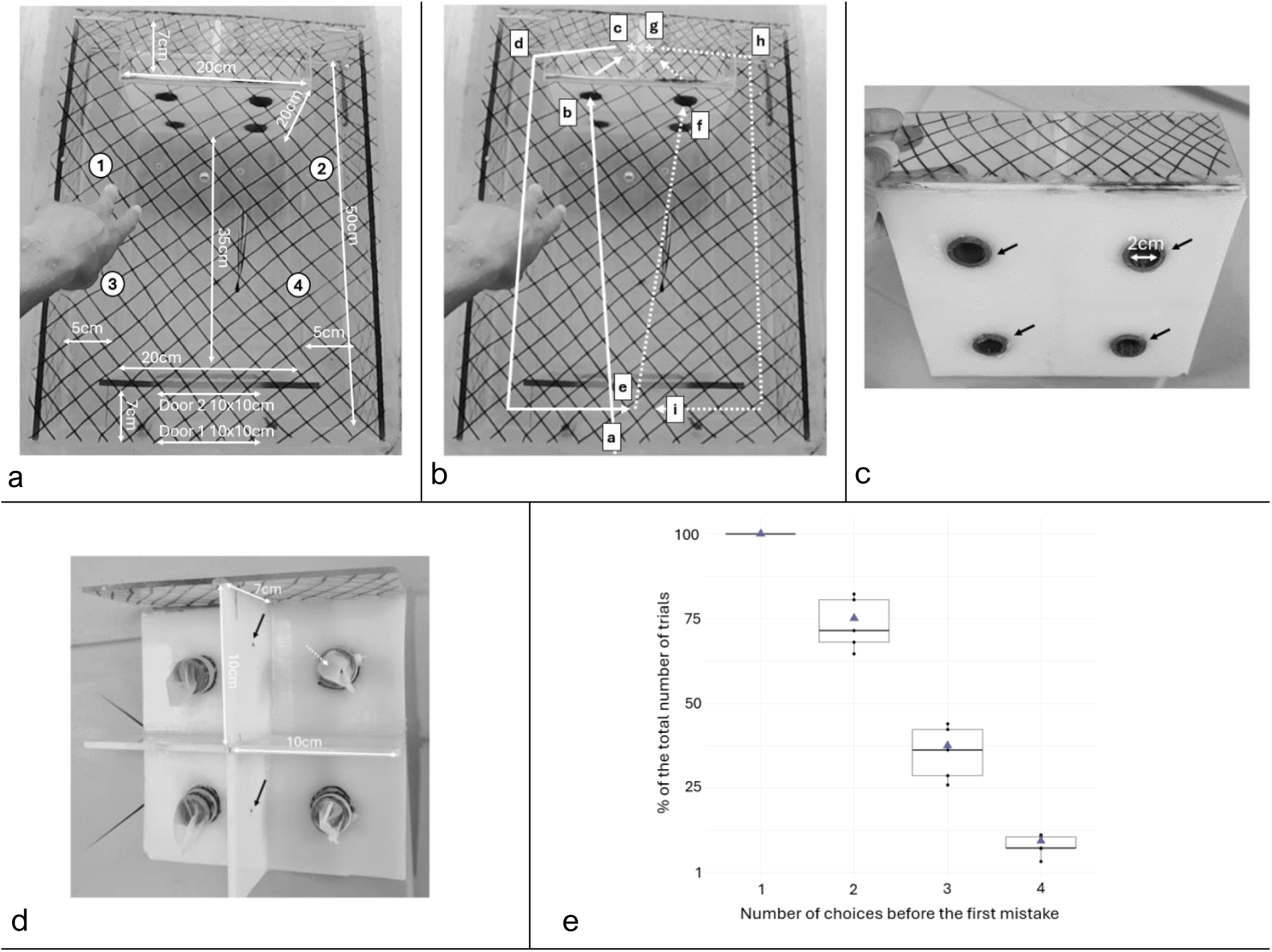
Working memory box, the movable testing arena. **a. Arena design.** The main door, “Door 1,” was manually operated by the experimenter, and cleaners entered the arena through this door to begin the trial. Swimming through the always open “Door 2” gave access to the central part of the arena and hence access to the plate with four pipes to access food behind. Corridors 1, 2, 3, and 4 lead from the respective compartments behind the plate back to the space between doors 1 and 2, allowing the fish to make a new run. Corridors 3 and 4 are positioned beneath 1 and 2, respectively. **b. Movements of cleaners in the arena.** An example in which a cleaner first chooses to visit compartment 1 and then compartment 2. Letters on arrows show the sequence of movements. The cleaner eats food items (*) at stages c and g. **c. Front view of the compartment plate.** Black arrows indicate the entrances to the four compartments. **d. Rear view of the compartment plate.** A food item was placed on the internal wall of each compartment (black arrows), and the white-dashed arrow indicates the rubber curtain. **e. Percentages of the number of trials in which cleaners succeeded in making 1, 2, 3, or 4 correct choices in a row.** The y-axis is the percentage of the number of trials, calculated on the usable trials only. The x-axis is the number of correct choices cleaners made before their first mistake. Because the first choice can never be a mistake, they succeeded in 100% of trials. Triangles indicate the random mean of success.

We provided extensive training over 15 days to habituate the cleaners to the complexity of the experimental setup. The training was conducted in intermediary steps: (a) picking a food item placed in the center of a dark circle drawn on a white plate; (b) replacing the dark circle with a piece of pipe; (c) inserting the pipe through a plate to create an entrance, leading the cleaner through it with food as a reward; (d) adding a piece of latex glove to the prototype; (e) utilizing the final compartment plate with food items positioned at their final locations (preventing the cleaner from changing compartments from the rear). Several reward plates were used to guide the fish through the box, particularly through the corridors. When the cleaner exhibited fear of the setup, the box was left in the aquarium with reward plates scattered around to encourage inspection. This process was repeated until the cleaner swam comfortably and swiftly through the corridors and through the entire box.

Trials started after subjects were habituated to additional elements of the experimental setup, such as swimming through doors, familiarizing themselves with different swimming routes within the box, and adapting to the box’s movement: we had a single box placed inside the aquarium right before the trial took place, which caused significant water displacement.

#### Data

Data collection was challenging. Some trials could not be completed, either due to external events causing behavioral disturbances, or because cleaners spent too much time in the corridors, or because the cleaner attempted to bypass the task by moving around the arena or enter the corridor from the wrong side to access food. Since we wanted to see how many choices cleaners could make correctly before the first mistake, we still considered a trial where the cleaner caused the experimenter to stop the trial on the third choice to see whether they could make two correct choices in a row. Ultimately, data from five cleaners were analyzed, with a variable number of usable trials for each individual.

##### Statistical analyses

We compared the probabilities of correctly selecting 2, 3, or 4 compartments consecutively by chance with the actual performance of cleaners. As it stands, if decisions are made by chance, the first choice correct is 100% probability, two choices correct in a row is 75% probability (3/4 compartments still contain food), three choices correct in a row is 37.5% probability (75% x 50%), and having all four choices correct has a probability of 9.375% (75% x 50% x 25%). For each cleaner, we made the sum of the first 2, first 3 and all 4 correct choices and incorrect choices over all its trials. We used a Wilcoxon signed-rank test to evaluate whether subjects consistently performed above chance levels.

#### Results

Cleaners failed to perform 2, 3, or 4 correct choices in a row above chance levels (Wilcoxon signed rank exact tests, 2 choices: V = 6 and p-val = 0.81, 3 choices: V = 5 and p-val = 0.63, 4 choices: V = 3 and p-val = 0.31, Fig. 2e). While the power of the test was very low, we note that observed means almost exactly match the predicted values, and not a single individual performed at clearly higher levels (Fig. 2e).

### 3. Experiment 3, two plates experiment

#### Methods

18 adult females, caught between 18. - 20.9.21, were tested on 13 days from 23.10. – 4.11.21. Three females did not properly participate as they were apparently scared by the procedure, leaving us with 15 subjects. We conducted 200 trials over 13 days. The number of trials per day varied between 5 and 20. Day 1 (trials 1-5) was viewed as training (methods described below as for day 3 onwards) and data were not analyzed. On day 2 (trials 6-15), we briefly tried a different method that we therefore analyzed separately from the large data set (see below). For the large data set, we divided the aquarium into three compartments using transparent barriers (Fig. 3a). Individual cleaners were initially in one of the side compartments. We then allowed them to eat a food item (mashed prawn) from the back of one plate in the middle compartment. The plate was either grey with a red pattern on both sides, or white with a green and yellow pattern on both sides (Fig. 3a). The plate was then removed, and after a waiting period of 5 to 10 seconds, the cleaner was presented with both plate types in the other side compartment, so that one was identical to the initial plate. On day 2, we had the two plates placed and visible to the cleaners before we introduced the first plate in the middle compartment. While this reduced the hectic to keep the time interval between first eating and second choice within 5-10s, we gave up on the design because the cleaner fish could obviously see that a total of three plates is involved, so that there would be no reason to consider the two identical plates as a unique one.

**Figure 3:**
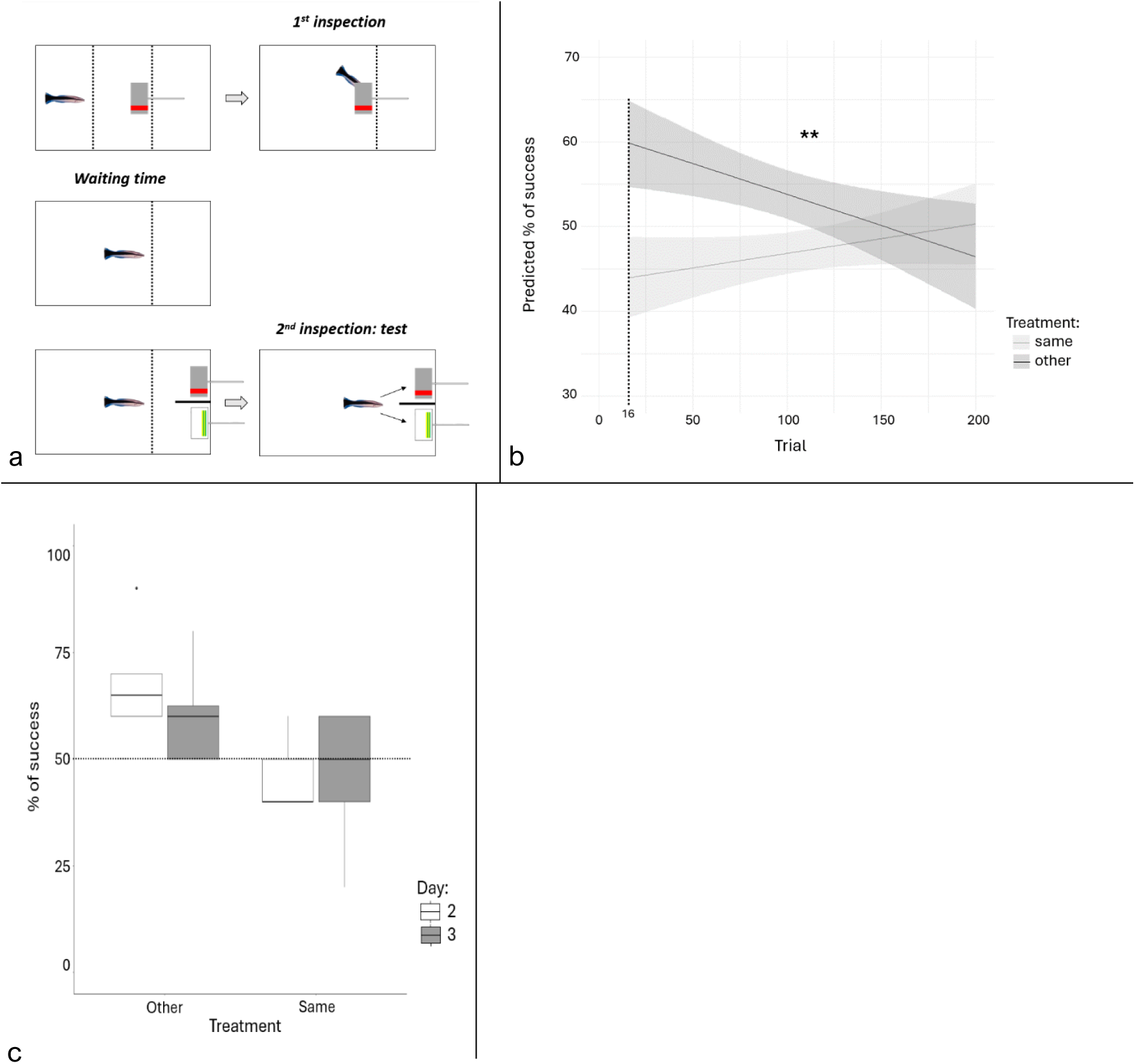
Two plates experiments. **a. The structure of a trial.** Dotted lines represent see-through barriers. Each trial involved two inspections, with the cleaner expected to make a choice during the second inspection based on the plate presented during the first inspection and the assigned treatment. The waiting time between inspections ranged from 5 to 10 seconds. **b. Different effects of time on the proportion of success depending on the treatment.** The y-axis is the predicted proportion (in percentages) of success, the x-axis shows the trial number, and the colors represent the treatments. Marginal effects plot showing the regression line and the 95% CI. In the “same” treatment, cleaners had to choose the plate that was the same as the one from the first inspection during the second inspection, while it was the other in the “other” treatment. The stars represent the significance of the interaction in the model. Significance codes: 0 ‘***’ 0.001 ‘**’ 0.01 ‘*’ 0.05 ‘.’ 0.1 ‘ ‘ 1. **c. Percentage of correct choices.** Boxplot showing the overall percentage of success (y-axis) of cleaners during the task, in both treatments (x-axis) and during day 2 and day 3. The dashed horizontal line indicates the random threshold of 50%.

Half of the cleaners were assigned to the “same” treatment, where they would find a food item behind the identical plate, while the other half were assigned to the “other” treatment, where the food item was placed behind the different plate.

The plate presented during the first presentation was counterbalanced, and no more than two consecutive trials featured the same plate. During the actual trial, the position of the plates was also counterbalanced across trials. Cleaners were allowed to inspect both plates, regardless of their first choice, as this approach has proven effective in facilitating learning in associative learning tasks. We scored a trial as ‘correct’ if the cleaner approached the rewarding plate first.

##### Statistical analyses

The main statistical analysis uses the data from trial 16-200. Data from day 2 (trials 6-15) were analyzed separately.

We performed a binomial generalized linear mixed-effects model, using the trial outcome (0: failure, 1: success) as the response variable, with treatment (“same” or “other”), trial number (from 16 to 200), and their interaction as explanatory variables. The identity of the cleaner was included as a random factor. Model assumptions were checked graphically using the diagnostic tools available in the *DHARMa* package (Hartig, 2022) in R.

Post-hoc analyses were conducted using the least-square means method from the *emmeans* package (Lenth, 2023) and simple slope effect calculations from the *reghelper* package (Hughes & Beiner, 2023) in R. The same statistical methods were applied to analyze data from day 2 separately, allowing for comparison between days 2 and 3.

#### Results

We found a significant interaction between the treatment and time (Fig. 3b). In the “same” treatment, the success rate of cleaners increased by 6.3% over time, but it was insignificant (Simple slope effect, slope = 1.001, t-value = 0.132). In the “other” treatment, it decreased by 13.4% (Simple slope effect, slope = 0.997, t-value = 0.008).

Overall, both treatments brought success rates close but significantly different from the 50% chance (Least-square means, same: predicted p of success = 47%, z-ratio = 2.41, p-val = 0.03 and other: predicted p of success = 53.6%, z-ratio = −2.4, p-val = 0.03).

During day 2 only, the two treatments were significantly different from each other as well (least-square means, other: predicted p or success = 67.5%, SE = 0.052 and same: predicted p of success = 44.4%, SE = 0.052, z-ratio = 2.99, p-val = 0.003) but only “other” treatment differed from chance (Least-square means, other: z-ratio = 3.06, p-val = 0.004 and same: z-ratio = - 1.05, p-val = 0.293).

The comparison between day 2 and day 3 did not indicate any difference (Type II Wald chisquare tests, day: chisq = 0.304, df = 1, p-val = 0.581 and treatment:day: chisq = 0.690, df = 1, p-val = 0.406, Fig. 3c).

### 4. Experiment 4, dynamic two plates experiment

#### Methods

23 cleaners (12 females, 11 males) that had been caught between 22.1. – 5.2.22. were tested in two groups, between 2. – 15.3.22. Subjects had been exposed to a foraging task before, so they were familiar with barriers and making choices at the onset of trials.

This experiment was very similar in design to experiment 3. The main difference was that the first presented plate was not placed into the middle compartment and then taken out of the aquarium to be replaced by a look-alike. Instead, both plates were present from the beginning in the rear compartment. Then, during the first inspection, one plate was moved towards the middle compartment and turned such that the food item on its back became accessible to the cleaner by putting its mouth through a hole pierced in the Plexiglas see-through separation. The plate was then turned and placed back to its initial position. The door in the middle of the separation was then lifted, and the cleaner had to inspect the back of the other plate in order to obtain a second food item (see the steps illustrated in Fig. 4a). Note that there were not two treatments as in experiment 3, the correct plate was always the one that was different from the first inspection.

**Figure 4:**
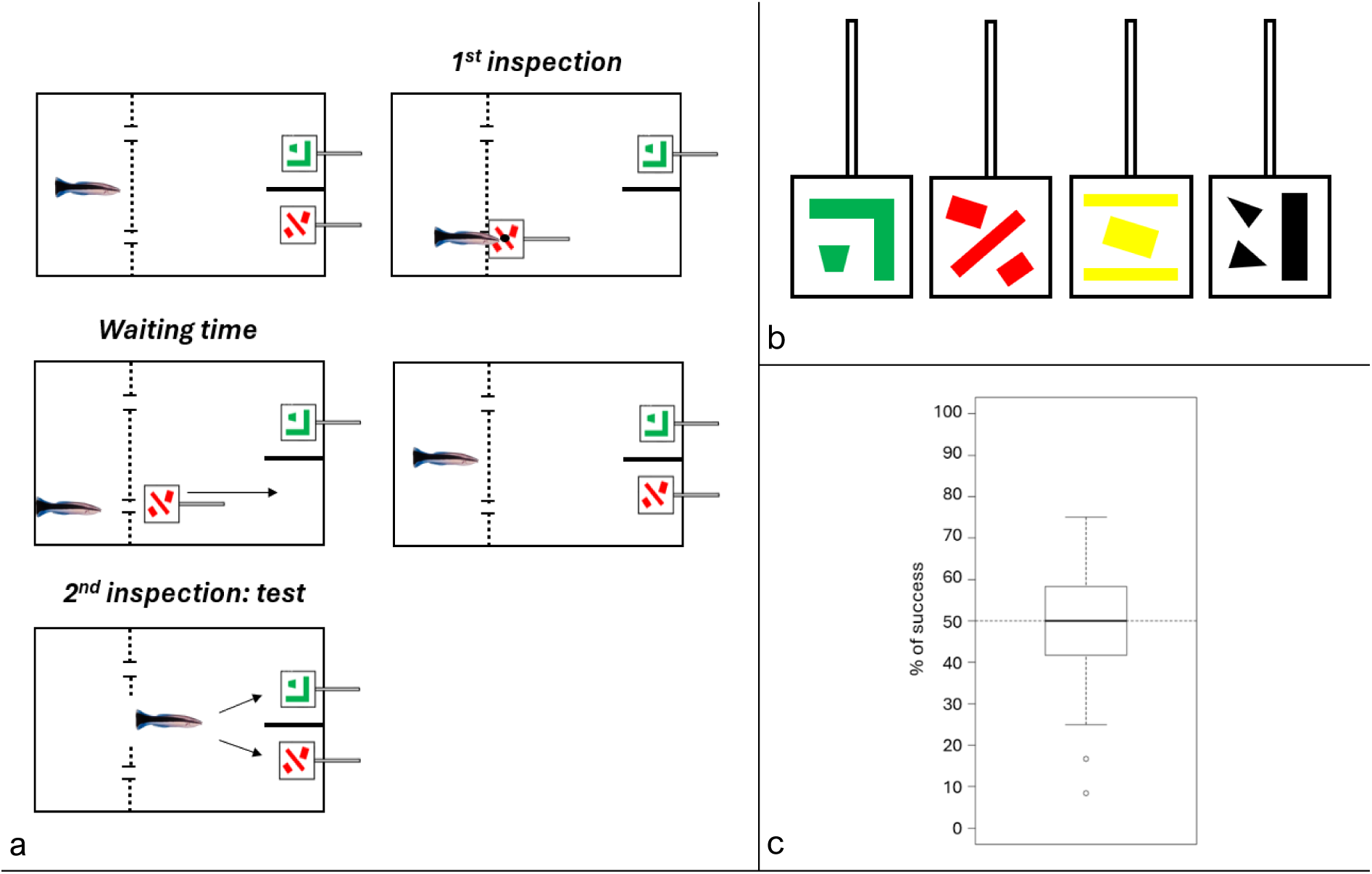
Experimental design. **a. Representation of a trial.** Dotted lines represent the see-through barrier, and the interruptions indicate two holes. For each trial, we inserted the initial plate in the water, turned it around while moving toward the barrier to expose the food item hidden in the back (black spot). After allowing the cleaner to eat, we returned the plate to the back of the aquarium. Following a waiting period of under 10 seconds, the barrier door was opened, and we noted the choice of the cleaner. **b. The four different plates.** Four different plates, each 5cm x 5cm with colored patterns (red, yellow, green, or black), were organized into six pairs. **c. Percentage of correct choices.** Boxplot showing the overall percentage of success (y-axis) of cleaners during the task. The dashed horizontal line indicates the random threshold of 50%.

Another difference to experiment 3 was that we used four plates in total that differed in color and pattern (Fig. 4b). All six possible combinations were presented, each twice per day for a total of 12 trials per fish per day. We counterbalanced positions and which plate became accessible across trials. The idea behind using four plates was to increase complexity if cleaners solve the task with two plates by adding a third and eventually a fourth plate, and varying the number of plates that a cleaner gets access to before the barrier is lifted. As we had some trials on the first day in which cleaners did not participate (eating through the hole appeared to be confusing at first), we analyzed only days 2-4.

We measured the time interval between the end of the first plate inspection and the moment the barrier door was opened for the second inspection, aiming to stay within a 5-10s interval. The choice of cleaners was noted. If it selected the same plate as in the first inspection, the trial was considered a failure, and both plates were removed before the cleaner could access the food item on the other plate.

##### Statistical analyses

Trials where the time delay between inspections was greater than 10 seconds were not considered as those were outside the limit.

Then, to account for variables that were unavoidable during data collection (e.g., the experimenter, the subject’s sex, the specific pair of plates and the trial number), we employed a binomial generalized linear mixed-effects model. The choice of the trial (0: failure, 1: success) was the response variable, and all variables mentioned were treated as covariates. The identity of the cleaners was included as a random factor. After none of the variables proved significant, we employed a null model to test the significance of the overall success rate.

For all models, we compared two versions using analysis of variance: one model with the cleaner’s identity as a random factor for both the intercept and slope with respect to time (trial number), and the other model with cleaners’ identity as a random factor for the intercept only. The simpler models were retained, as they showed no significant difference and had slightly lower Akaike Information Criterion (AIC) values.

We graphically checked the model assumptions using the diagnostic tools from the *DHARMa* package (Hartig, 2022) in R. The significance of the variables was tested using Type II Wald Chi-square tests, and post-hoc analyses were conducted using the least-square means method from the *emmeans* package (Lenth, 2023).

#### Results

Focusing only on trials in which the delay was between 5-10s, we could use a total of 1038 trials out of 1102 (94.2 %). None of the covariates we studied had a significant effect on the choice of cleaners (Type II Wald chisquare tests on binomial glmer, experimenter: chisq = 0.028, df = 1, p-val = 0.868; sex of fish: chisq = 0.117, df = 1, p-val = 0.733; pair: chisq = 3.034, df = 5, p-val = 0.695; trial: chisq = 0.504, df =1, p-val =0.478).

The null model indicated that cleaners did not visit the correct plate above chance (Least-square means, predicted mean = 52.3%, SE = 0.016, z-ratio = 1.489, p-val = 0.136, Fig. 4c).

## 5. Discussion & Conclusion

This study comprises four novel experimental setups that may be indicative or explicitly test for WM in cleaners. Overall, the performance of cleaners was modest, even in the first two tasks that could, in principle, be solved without WM. Significant results were weak and inconsistent, like an interaction with the number of trials in experiment 3 in which the initial effect disappeared over time. The absence of clear positive evidence in any task made it unnecessary to conduct further testing aimed at ruling out alternative problem-solving mechanisms, like the development of a feeding sequence routine in experiment 1 or the use of familiarity in experiment 2 (i.e. the ability to react to a stimulus without explicit recollection of the information associated to it; see the difference between familiarity and recollection in Yonelinas, 2002). Similarly, the lack of positive evidence also made it unnecessary to test for sensitivity to disturbance or storage capacity of WM (e.g., number of items that can be held and manipulated simultaneously), two key aspects often associated with WM capacity and largely overlooked in the literature. We recognize that the absence of evidence cannot be perfect proof for evidence of absence, but consistent results across the four different experiments make us skeptical about the existence of WM in cleaners. Previous laboratory experiments with cleaners highlighted the use of strategies that capitalized on previous knowledge and experiences (Binning et al., 2017; Bshary & Grutter, 2006; Grutter, 2004; Pinto et al., 2011a). Because the deployment of such strategies seems to require agile behavioral adjustments, relying on executive control and adaptability, we had expected WM to be recruited (Baddeley & Hitch, 2019; Cowan, 1995; Diamond, 2013).

We provide detailed discussions of each experiment in the supplementary material, including details of earlier setups that were confusing for cleaners and hence, were not fully carried out.

### Link to ecology

We strived to design experiments that could reflect the ecology of cleaner fish, with the goal of engaging their WM abilities, if such abilities exist. For example, when cleaners clean their clients, they inspect different surface locations (*e.g.,* mouth, gills, tail). For the sake of efficiency, it would be expected that cleaners remember and hence, avoid revisiting previously inspected locations (i.e., they optimize their foraging strategy). Avoiding previous locations should reflect the ability to store and update information, a key signature of WM. Our “windows experiment” and the “movable arena experiment” aimed to elicit this ability. We found that cleaners were more efficient when windows were overlaid on an image of a client fish (here, a butterflyfish) *versus* overlaid on a plain white square plate. The picture of a client fish was intended to give additional ecological relevance to the experimental setup. While we found significant effects or a tendency that plate design matters, we currently cannot rule out that simple differences in coloring (rather than fish *versus* plate) may have affected the behavior of cleaners. In any case, the more important observation was that the presence of windows strongly reduces the cleaners’ foraging efficiency, independently of plate type.

The other two experiments, where we used pairs of plates, were also intended to simulate cleaners’ ecology. Because cleaners have multiple repeated interactions per day (Grutter, 1996a), we hypothesized that they would likely remember recent clients to avoid inspecting the same individual again, thereby optimizing their foraging behavior. Indeed, a previous study found that cleaners could adapt their foraging choices based on information they acquired up to several minutes before, avoiding “client plates” that they had inspected within the time windows (2.5 to 12.5 min) set by the experimenter (Salwiczek & Bshary, 2011). In other words, cleaners could remember “*when* they interacted with *what* after a single event” (Salwiczek & Bshary, 2011). Although the possession of something akin to episodic-like memory would account for these results (Pause et al., 2013; Tulving, 2002), it seemed reasonable to expect that their ecology selects cleaners to succeed also when the time intervals separating acquisition and retrieval is shorter, which would require WM. Lack of success in this new condition reinforces the idea that time is a critical factor for eliciting WM and that the separation between LTM and WM is a valid one, even if one is to accept that WM could operate with information reactivated from LTM (Cowan, 2008, 2019; Norris, 2017).

### The lack of WM in cleaners, and other ectotherms?

We did not anticipate that cleaners would perform so poorly in all four experimental tasks, but the lack of positive evidence for WM fits the apparent absence of a general intelligence factor *g* reported for this species (Aellen et al., 2022). In mammals, individual performance in WM tasks is a reliable predictor of performance in multiple other cognitive tasks, indicating that the presence of WM is a precondition for the presence of the *g* factor (Ackerman et al., 2002; discussed and reviewed in Conway et al., 2003; Engle et al., 1999; Shipstead et al., 2014). Consequently, we did not expect that the performance of cleaners would be comparable to that of large-brained endotherm species (e.g., primates or corvids). Another potential difference could have laid on the sensitivity to distraction, with cleaners being possibly more sensitive have also expected that cleaners would be more easily distracted (discussed in Carruthers, 2013 and Manrique & Walker, 2017), but this hypothesis needs specific testing using paradigms where the fish show initial success in the absence of disturbance. Nevertheless, failing to find any significant evidence for WM in cleaners, within ecological validity and on a low level, came as a surprise. As it stands, species differences in WM capacities can in principle be quantitative, as for example between humans and chimpanzees (Manrique et al., 2024). Our experiments 3 and 4 were designed such that cleaners could have performed at a low level (above chance for the second choice in the movable arena experiment, above chance as long as only two plates are present in the dynamic two plates experiment) but not when the required information increases (like for the 3^rd^ and 4^th^ choice in the movable arena experiment, or when confronted with 3 or more choices in the dynamic two plates experiment). Success with a single choice being indicative of a WM of size 1, we had expected that such a small capacity would be present in this species given their proficient problem solving, yet data proved us wrong.

A question arising from our study is whether the absence of WM is specific to cleaners, or is a key difference that separates the cognition of ectotherms and endotherms (Jerison, 1969). The consensus in the literature is that WM is a system that keeps recently acquired information in an active state, allowing its combination with knowledge previously acquired, stored in the long-term memory and reactivated, to assist prospective adaptive action (e.g., Baddeley & Hitch, 1974; Cowan, 1995, 2019). With this definition in mind, we consider the extant literature regarding WM in fish to be very incipient. Studies to-date have investigated the natural alternation rate by zebrafish in a Y-maze during a free exploration task (Fontana et al., 2021) or showed zebrafish succeeding in a delayed match-to-sample experiment (Bloch et al., 2019). Furthermore, both cichlids and guppies selected for a bigger telencephalon size showed significant, though modest, success in the Piagetian stage 4 of object permanence (i.e., recovering a hidden object, Piaget & Cook, 1952), which the authors directly associated with WM (Guadagno & Triki, 2024; Triki et al., 2023). These are all interesting results that can be compared to the performance of endotherms. However, these paradigms may not test for WM. Maze studies to measure WM have been criticized for not considering whether the time interval separation between acquisition and retrieval is within a range compatible with WM (Carruthers, 2013). In addition, the manipulative and problem-solving aspect of WM is barely detectable in the task, especially when it is about purely explorative behavior, as in Fontana et al. (2021). Match-to-sample tasks are visual tasks where processes like familiarity could satisfactorily account for the observed positive results, without invoking higher-level executive control (e.g. Aggleton et al., 1986; Cowan, 2019; Yonelinas, 2002). That is why they are often considered “recognition tasks” (Bachevalier & Mishkin, 1986; Zola et al., 2000). Regarding experiments on object permanence, we argue that only later stages (i.e., Piagetian stages 5 and 6) can reliably be linked to WM, as the combination and manipulation of information are absent in previous stages. In Piagetian stage 4 (single visible displacement task, Piaget & Cook, 1952), the subject must only retrieve a hidden object. Therefore, we would argue that there is no necessity for the combination or manipulation of previously acquired information. In contrast, later stages (5 and 6, with invisible displacement, Piaget & Cook, 1952) almost certainly engage WM, as they require varying degrees of mental computation to track the trajectory of an unseen moving object.

Thus, to answer the question on the presence/absence of WM in ectotherm vertebrates, we need to conduct experiments that are explicitly designed to test for this executive function. Importantly, we are confident that performance above chance in experiments 3 and 4, in combination with a lowered performance when a disturbance before the choice is introduced, would provide conclusive evidence for the presence of WM (Carruthers, 2013; Cowan et al., 2024). These paradigms can be applied to other species, allowing for cross-species comparisons.

In conclusion, our comprehensive series of experiments yields no evidence for WM abilities in cleaners, which contrasts with evidence for WM in endotherm vertebrates, including corvids and psittacids (Pepperberg & Funk, 1990; Zucca et al., 2007), mice (Kolata et al., 2005), rats (Bratch et al., 2016) and chimpanzees (e.g., Völter et al., 2019, debates in Manrique et al., 2024; Read et al., 2022). This contrasts with research on the other two main executive functions (EFs) – cognitive flexibility (Aellen et al., 2022; Parker et al., 2012) and inhibitory control (Aellen et al., 2021; Lucon-Xiccato & Bisazza, 2017; Sovrano et al., 2018) – for which evidence has been found in fish, including cleaner wrasse (Aellen et al., 2022). Taken together, the results raise the intriguing possibility that rather than all executive functions, WM and the associated *g* factor warrant an enlarged and reorganized brain as found in endotherms but not in ectotherms (Jerison, 1969).

## Acknowledgements

The staff of the Richard B. Gump research station in Mo’orea, French Polynesia. S. Duc, E. de Sousa, and R. Bonvini for helping during data collection, Prof. K. Zuberbüler for his help in conceptualizing the movable arena, Dr. Radu Slobodeanu for statistical advice, and Dr. C. S. Clements for his help in improving the writing. Funding was provided by the Swiss Science Foundation, grant to R.B number 310030_192673/1.

